# Hypoxia Regulation of *ndrgs*

**DOI:** 10.1101/2020.12.16.422782

**Authors:** Nguyet Le, Timothy Hufford, Rachel Brewster

## Abstract

Many organisms rely on oxygen to generate energy in the form of adenosine triphosphate (ATP). During severe hypoxia, the production of ATP decreases due to diminished activity of the electron transport chain, leading to cell damage or death. Conversely, excessive oxygen causes oxidative stress that is equally damaging to cells. To mitigate pathological outcomes, organisms have evolved mechanisms to adapt to fluctuations in oxygen levels. Zebrafish embryos are remarkably hypoxia-tolerant, surviving anoxia (zero oxygen) for hours in a hypometabolic, energy-conserving state. To begin to unravel underlying mechanisms, we analyze here the distribution and hypoxia-dependent regulation of members of the N-myc Downstream Regulated Gene (Ndrg) family, Ndrg 1-4. These genes have primarily been studied in cancer cells, and hence little is understood about their normal function. We show here using *in situ* hybridization that, under normoxic conditions, *ndrgs* are expressed in metabolically-demanding organs of the zebrafish embryo, such as the brain, kidney, and heart. Following exposure of embryos to different severity and durations of hypoxia, we observed that *ndrgs* are differentially regulated and that *ndrg1a* is the most responsive member of this family, with nine-fold upregulation following prolonged anoxia. We further show that this treatment resulted in *de novo* expression of *ndrg1a* in tissues where it is not observed under normoxia, such as head vasculature, the inner ear, and somites. These findings provide an entry point into understanding the role of this conserved gene family in hypoxia adaptation of normal cells.

## INTRODUCTION

The air in earth’s atmosphere is made up of approximately 21 percent oxygen (O_2_). Aerobic organisms use this environmental O_2_ to produce ATP during oxidative phosphorylation. Hence fluctuations in O_2_ levels, either up or down, can have very detrimental outcomes for aerobic organisms. Severe hypoxia causes a decrease in ATP production due to diminished activity of the electron transport chain. Given that ATP fuels energy-demanding processes in the cell, its reduction can lead to cellular damage or death [1–4]. Thus, hypoxia, hypoxemia or ischemia, is a contributing cause to many disease states in humans, including pulmonary vascular disease, acute kidney injury, neurodegenerative disease, and stroke [5–10]. Conversely, excessive O_2_ is equally, if not more harmful as it causes oxidative stress due to reactive oxygen species production that is damaging to macromolecules, including lipids, proteins, and nucleic acid [11–13].

To mitigate these adverse consequences, aerobic organisms have evolved mechanisms to adapt to low O_2_ levels and maintain homeostasis. Such adaptations optimize access to O_2_ by increasing red blood cell count and angiogenesis and altering energy metabolism in part by switching from oxidative phosphorylation to glycolysis [14, 15]. In addition, cells conserve energy when exposed to chronic and severe hypoxia by reducing their metabolic rate. The latter is accomplished via suppression or arrest of energetically-demanding processes such as cell division, transcription and translation and down-regulating the activity of the sodium-potassium ATPase pump [16–23]. While metabolic suppression has primarily been studied in organisms considered anoxia-tolerant, including painted turtles, crucian carp, naked mole rats and hibernating ground squirrels [16, 24], it is likely to also be utilized in other organisms as well, albeit to a lesser extent. Zebrafish *(Danio rerio)* embryos maintain homeostasis under anoxia (zero O_2_) by entering into a hypometabolic state characterized by reversible developmental arrest, which enables them to survive for up to 50 hours [25, 26]. This protective response is developmentally regulated, with older embryos being less tolerant to anoxia [26].

Despite the necessity to conserve energy via suppression of transcription and translation, genes that are vital for the hypoxia response are in fact transcriptionally up-regulated under hypoxia [27–30]. Such upregulation is mediated by several transcription factors, the best studied of which is the Hypoxia-Inducible Factor-1α (HIF-1α) [31–33]. Under normoxic conditions (normoxia), the HIF-1α subunit is hydroxylated by prolyl hydroxylase domain protein 2 (PHD2), marking it for degradation by the von Hippel-Lindau protein (VHL). However, when O_2_ levels are reduced, PHD2 activity is inhibited and stabilized HIF-1α binds to the HIF-1β subunit and translocates to the nucleus to regulate transcription. Upon entry into the nucleus, HIF-α/β heterodimers bind the hypoxia-response element (HRE). Even though this sequence is abundant in the genome, fewer than 1% of potential HRE sites are bound by the HIF complex under hypoxia, suggesting the existence of another layer of regulation [34–37]. HIFs directly activate genes that mediate metabolic reprogramming from oxidative phosphorylation to glycolysis [38, 39] and genes that increase the available O_2_ supply, such as *EPO, VEGF* and its receptors [40]. Other HIF targets are implicated in autophagy, apoptosis, redox homeostasis, inflammation and immunity, stemness and self-renewal, metastasis and invasion [35, 39, 41, 42]. In addition to HIFs, several other transcription factors are known to influence the hypoxia response, including CREB, Myc, NF-kB, and STATs, which engage in cross-regulatory interactions with HIFs [32].

Members of the N-myc downstream regulated gene (NDRGs) family are also hypoxia-responsive. The mammalian family consists of 4 members, *NDRG1-4,* while the zebrafish genome with its third round of genome duplication, encodes 6 paralogues, *ndrg1a, 1b, 2, 3a, 3b,* and *4* [43]. NDRGs are highly conserved across metazoans and the sequence homology is in fact greater for specific members of the family across different species (> 80%) than between NDRG family members of the same species, which share ~ 57–65% amino acid identity [44]. NDRGs belong to the α/β-hydrolase family, however they are enzymatically inactive, lacking a critical catalytic triad [45]. NDRG1 (formerly known as Drg1, Cap43, Rit42, RTP, and PROXY-1) contains three tandem repeats (GTRSRSHTSE) near its C-terminal and a phosphopantetheine sequence, which are two unique features that make it distinct from other NDRG family members. NDRG1 is thought to function as a tumor suppressor [46]. However, the absence of cancer resultant from germline mutations in humans [47] and targeted knockout in mice [48], suggests that *NDRG1* may rather be involved in cancer progression (metastasis) rather than initiation [49–54]. Human NDRG1 interacts with numerous other proteins in human cancer and other cell lines, including actin, Clathrin and associated protein AP-1 and AP-2, Caveolin-1, Kinesin, LAMP1, Rab4, and 26S proteasome components [49, 51, 55, 56], consistent with a possible role in regulating vesicle trafficking [56, 57]. *NDRG1* and *NDRG2* transcript levels increase under hypoxia, as these genes have HIF-1α binding sites (hypoxia-response elements or HREs) in their promoters [58–61]. However, NDRG regulation under hypoxic conditions is complex and does not depend solely on HIF-1α, as several other transcription factors [62, 63] and *NDRG1* long non-coding RNA itself [64, 65] have also been implicated. *NDRG4* is transcriptionally up-regulated under hypoxia in cancer cells, however in a TNF-α/NF-κB rather than a HIF-1α-dependent manner [66]. In contrast, NDRG3 is regulated post-translationally in hypoxic cancer cells, via lactate binding, which stabilizes the protein and promotes cell proliferation and angiogenesis [67]. These findings indicate that NDRGs are regulated at the transcriptional and post-translational levels in response to hypoxia and promote adaptation to low O_2_.

To date, NDRGs have mostly been studied in cancer cells and far less is therefore known about their normal function and regulation. While all members of this family can be regulated in response to fluctuations in O_2_ levels, it is unclear what range and duration of hypoxia they respond to. Lastly, even though the spatial distribution of *NDRG* family members has been analyzed in zebrafish [68, 69] and frog *(Xenopus laevis and tropicalis)* embryos [43, 70, 71] and mammals [72–80], it is unclear whether their spatial distribution changes under low O_2_. We report here on the spatial distribution of members of the zebrafish Ndrg family and their regulation in response to hypoxia.

## MATERIALS AND METHODS

### Zebrafish

Zebrafish *(Danio rerio)* were raised and housed at 27°C on a 14/10 hour light/dark cycle. Zebrafish used in this study were the wild-type AB strain. Embryos were obtained by breeding male/female pairs. Maintenance of zebrafish and experimental procedures on larvae and adult zebrafish were performed in accordance with the protocol approved by the Institutional Animal Care and Use Committee (IACUC) at the University of Maryland Baltimore County. Zebrafish embryos (raised in normoxia) were staged and sorted according to Kimmel *et al.* 1995 [81]. See Supplementary Figure 2 for staging of anoxia-treated embryos.

### Hypoxia & anoxia treatments

For whole-mount *in situ* hybridization (WISH), 24-hour post fertilization (24 hpf) zebrafish embryos were dechorionated and then placed in 100 mm petri dish (CellTreat, Pepperell, MA, USA, Cat# 229663) containing 0% or 3% O_2_ system water in an O_2_ control glove box (Plas-Labs, Lansing, MI, USA model # 856-HYPO) set at 0% or 3% O_2_ and 27°C. Following 8 h of treatment, embryos were placed into 1.5mL microcentrifuge tubes (20 embryos/tube) with excess water removed. Embryos in each microcentrifuge tube were removed from the chamber and fixed in 4% paraformaldehyde (PFA) in phosphate buffer saline (PBS) (Thermo Scientific, Waltham, MA, USA Cat# J19943-K2) at 4°C overnight. Fixed embryos were rinsed in absolute methanol (Thermo Fisher Scientific, Waltham, MA, USA, CAS# 67-56-1) for 10 min at room temperature and stored in absolute methanol at −20° C.

For Real-time qPCR, stage-matched control embryos were placed in 100 mm petri dishes containing 0% or 3% O_2_ system water, placed in the O_2_ control glove box set at 0% or 3% O_2_ and 27°C. Following 4h and 8h of treatment, single embryos were placed into 1.5mL microcentrifuge tubes (Stellar Scientific Ltd Co, Albuquerque, NM, USA, Cat# T17-100) with excess water removed. Single embryos in the microcentrifuge tubes were taken out of the chamber, flash frozen in liquid nitrogen, and stored at −80°C for total RNA extraction. Embryos raised under normoxic conditions were used as stage-matched controls for hypoxia (27hpf for 4h and 30.5hpf for 8h) and anoxia (26hpf for 4h and 27hpf for 8h) treatments.

The PCR template was cDNA synthesized from total RNA extracted from a combination of 6 hpf, 24 hpf and 48 hpf zebrafish embryos. RNA extraction was performed with the QuickRNA MicroPrep Kit (Zymo Research, Irvine, CA, USA Cat#R1051) and cDNA synthesis was carried out using the iScript cDNA Synthesis Kit (Bio-Rad, Hercules, CA, USA Cat# 1708890) according to the manufacturers’ instructions.

PCR reactions were prepared using 1 μl of diluted (1:5) cDNA as template in a total volume of 50 μl. Primer concentrations were 10 μM for each oligonucleotide. PCR-fragments were produced on a C1000 Thermal Cycler (Bio-Rad, Hercules, CA, USA Cat#1851148) using Phusion-Polymerase (Thermo Scientific, Vilnius, LT, F530S) (35 cycles and 57°C annealing temperature). PCR-fragments were gel-purified using Micro Bio-Spin P-30 Gel columns Tris Buffer (RNase-free)(Bio-Rad, Hercules, CA, USA Cat#7326250) and subsequently, 300-500 ng were used as template-DNA to synthesize antisense RNA probes by *in vitro* transcription using the mMESSAGE mMACHINE T7 Transcription Kit (Thermo Fisher Scientific, Carlsbad, CA, USA Cat#AM1344), incorporating digoxigenin (DIG)-UTP via a DIG-labelling kit (Roche, Mannheim, Germany Cat#11277073910).

To avoid amplification of regions of homology between *ndrg* members, all oligonucleotide primer pairs were designed against the 3’UTR of each gene, with the exception of *ndrg2* for which the primer pairs targeted the coding region (spanning exons 11-16) as well as the 3’UTR, as specified in Li *et al.* 2016 [68]. For antisense probes, a T7 promoter sequence **(5′- TAATACGACTCACTATAG-3′)** was added to the 5′ end of each reverse primer. The following primer sets were used to amplify cDNA for 35 cycles as follows:

*ndrg1a* forward: 5′-ACCAATCAGTTCTGACTGTGCTGC-3′

*ndrg1a* reverse: 5′- **TAATACGACTCACTATAG**CACTCCCAACATGGAAAACGCAGA-3′

*ndrg1b* forward: 5′- ACACGCCTCAGCAGTTTAATCTGG-3′

*ndrg1b* reverse: 5′- **TAATACGACTCACTATAG**CTCACTGAAGTCTTGCACAACCAG-3′*ndrg2* forward: 5′- ACAACACGTTCAAATGCCCG-3′

*ndrg2* reverse: 5′- **TAATACGACTCACTATAG**GGAAGACATGAGCTGGCTGT-3′

*ndrg3a* forward: 5′- GGTCTTCCAACTGGTTTGAGATGC-3′

*ndrg3a* reverse: 5′- **TAATACGACTCACTATAG**TGAGAACCAGTGGACAGTGACACT-3′

*ndrg3b* forward: 5′- GCCAGAGAGTGCTGGTCTAATGAA-3′

*ndrg3b* reverse: 5′- **TAATACGACTCACTATAG**CCGAGACATGCTAATCAGTAGCTC-3′*ndrg4* forward: 5′ - GACTTGCGTCAGGGATGATAACCT-3′

*ndrg4* reverse: 5′- **TAATACGACTCACTATAGG**AATGAGTGAGAGCAAGGGCCGAT-3′

### Whole-mount RNA *in situ* hybridization

Normoxic controls were fixed at desired stages (shield, 15-somites, 24 hpf, and 48 hpf) in 4% PFA in PBS (Thermo Scientific, Waltham, MA, USA Cat# J19943-K2) at 4°C overnight. Anoxia-exposed 24 hpf embryos were treated as described in the anoxia treatment section above. To prevent pigmentation from masking the WISH signal, embryos fixed after 24 hpf were incubated in 0.003% 1-phenyl-2-thiourea (PTU) (Sigma-Aldrich, Milwaukee, WI, USA Cat# P7629-100G) at 24 hpf until the time of fixation. Whole-mount single colorimetric RNA *in situ* hybridization (ISH) was performed on both normoxic and anoxic embryos using DIG labeled antisense probes according to the specifications published by Thisse and Thisse (2008). Briefly, embryos were rinsed in PBS (137mM NaCl, 2.7mM KCl, 8.8mM Na2HPO4)with 0.1% Tween-20 (Sigma-Aldrich, Milwaukee, WI, USA CAS# 9005-64-5). Proteinase K (10 μg/ml)(Sigma-Aldrich, Milwaukee, WI, USA CAS# 39450-01-6) treatment was performed for 10 min (24hpf), 12 min (27 hpf control and 24 hpf + anoxia treated), and 30 min (48 hpf) embryos. Embryos were then hybridized with DIG labeled antisense probes (*in situ* hybridization mix with dextran) at 70°C overnight. Following hybridization, excess probes were removed by washing embryos in a saline-sodium citrate (SSC) series (Thermo Fisher Scientific, Waltham, MA, USA, Cat# AM9763). For probe detection, alkaline phosphatase-conjugated antibody (Roche Diagnostics, Mannheim, Germany Cat# 11093274910) diluted (1:10,000) in pre-incubation (PI) buffer (PBS, 0.1% Tween 20, 2% sheep serum, 2mg/ml BSA (Thermo Fisher Scientific, Waltham, MA, USA, CAS# 9048-46-8) were added and incubated at 4°C overnight. 5-bromo-4-chloro-3-indolyl-phosphate (BCIP) (Roche, Mannheim, Germany Cat#11383221001) was used in conjunction with nitro blue tetrazolium (NBT) (Roche, Mannheim, Germany Cat#11383213001) for the colorimetric detection of alkaline phosphatase activity. When the color was optimal, the reaction was stopped by washing in PBS with 0.1% Tween-20 and rinsed in 4% PFA overnight.

### Vibratome sectioning, microscopy, and imaging

Following WISH for *ndrg1a,* embryos were embedded in 4% low melt agarose (IBI Scientific, Peosta, IA, USA, Cat# IB70050) in PBS (100 g/ml) and sectioned using a vibratome (Vibratome,1500) set at 40μm thickness. Cross sections were mounted on glass slides in 50% glycerol (Sigma-Aldrich, Milwaukee, WI, USA, Cat# G7757-1GA) under cover slips.

Zebrafish embryos were mounted for imaging from a lateral or dorsal view on slides on top of 4% w/v methylcellulose (Sigma Aldrich, Milwaukee, WI, USA, Cat# 274437-500G) in 1X E3 medium (5mM NaCl, 0.17mM KCl, 0.33mM CaCl2, and 0.33mM MgSO4)(for zebrafish embryo). Bright-field images were captured using an AxioCam HRc 503 CCD camera mounted on an Axioskop (Carl Zeiss, Oberkochen, Germany).

### Real-time quantitative PCR

RNA extractions and cDNA synthesis were carried out using single embryos collected at the appropriate developmental stage for the stage-match controls and immediately following treatment for anoxia- and hypoxia-treated embryos. RNA extractions were performed using the QuickRNA MicroPrep Kit (Zymo Research, Irvine, CA, USA Cat#R1051) cDNA was synthesized from total RNA using iScript cDNA Synthesis Kit (Bio-Rad, Hercules, CA, USA Cat# 1708890) according to the manufacturers’ instructions. The cDNA samples were diluted 1:10 with nuclease-free water (Life Technologies Corp., Austin, TX, USA, Cat# AM9937) Real-time quantitative PCR (qPCR) experiments were carried out with a CFX96 Touch Real-time qPCR Detection System (Bio-Rad, Hercules, CA, USA) using the SsoAdvanced Universal SYBR Green Supermix (Bio-Rad, Hercules, CA, USA Cat# 172-5271). Levels of mRNA of *ndrg1-4* were evaluated and *ef1a* was used as a reference gene. All oligonucleotide primer pairs included regions common to all spliced variants of each *ndrg* member, with the exception of *ndrg1b* and *ndrg2,* which only have one variant. The following primer sets were used to amplify cDNA for 30 cycles as follows:

*ndrg1a* forward: 5′-ATCATGCAGCACTTCGCTGT-3′

*ndrg1a* reverse: 5′-CAATAGCCATGCCGATCACA-3′

*ndrg1b* forward: 5′-CATGGGCTACATGCCCTCTG-3′

*ndrg1b* reverse: 5′-TGACCCGATGAACTGTGCTC-3′

*ndrg2* forward: 5′-AGCTGGAAAGAAAGTGCGAGA-3′

*ndrg2* reverse: 5′-TTTACGCCGTCCGCTTATGT-3′

*ndrg3a* forward: 5′-GGACTAGCAATCTTGTGGAC-3′

*ndrg3a* reverse: 5′-TCTCGATTCCGAGGTCTTGA-3′

*ndrg3b* forward: 5′-GTCAGGCTTGATGATGGATG-3′

*ndrg3b* reverse: 5′-CCCTCTCAAAGTCACATGAAGG-3′

*ndrg4* forward: 5′-AGCCAGCTATTCTGACCTAC-3′

*ndrg4* reverse: 5′-GATATCCTTGAGGCATCTGG-3′

*ef1a* forward: 5′-TACCCTCCTCTTGGTCGCTT-3′

*ef1a* reverse: 5′-TGGAACGGTGTGATTGAGGG-3′

Reactions were run in triplicate with 8 biological replicates, using 1 μl of diluted cDNA as template in a reaction volume of 20 μl. Primer concentrations were 10 μM for each oligonucleotide (Thermo Fisher Scientific, Waltham, MA, USA). Evaluation of results was performed in the CFX96 Touch RT-PCR Detection System program (Bio-Rad, Hercules, CA, USA) and using GraphPad Prism 8 software (Prism, San Diego, CA, USA).

## RESULTS

### The spatial distribution of the *ndrg* family during early development

The zebrafish genome encodes 6 homologs of *ndrg* family: *ndrg1a* (ENSDARG00000032849), *ndrg1b* (ENSDARG00000010420), *ndrg2* (ENSDARG00000011170), *ndrg3a* (ENSDARG00000013087), *ndrg3b*(ENSDARG00000010052), and *ndrg4* (ENSDARG00000103937). To characterize the members of the *ndrg* family, we began by determining their spatial distribution in early-stage zebrafish embryos, using whole-mount *in situ* hybridization (WISH). Since several members of the *ndrg* family contain large overlapping coding sequences (53-65% homology), riboprobes were designed that bind to non-conserved regions in the 3′UTR of all *ndrgs,* with the exception of *ndrg2.* Due to issues with the amplification of the *ndrg2* 3′UTR, we used instead a riboprobe complementary to the coding region and 3′UTR of this gene that has low homology with other *ndrg* members [68].

During zebrafish early development, *ndrg1a* expression is ubiquitous at the shield stage (Fig. 1A). During segmentation (15-somites), it becomes restricted to the neural crest, pronephric ducts (embryonic kidney) and ionocytes (also known as mucous cells) (Fig. 1B). At 24 hpf *ndrg1a* is also weakly expressed in the epiphysis, in addition to the pronephric duct, ionocytes and caudal vein (Fig. 1C,D) and by 48 hpf, *ndrg1a* is observed in corpuscles of Stannius (endocrine glands in the kidney), liver, intestinal bulb, retina, and other brain regions (Fig. 1E,F). The expression of *ndrg1b* is very dynamic. At shield stage it is ubiquitously expressed (Fig. 1G), while by 15-somites and 24 hpf, it is no longer detected (Fig. 1I,J). In contrast, at 48 hpf, *ndrg1b* is strongly expressed in the retina (Fig. 1K,L). The expression of *ndrg2* is ubiquitous at shield stage (Fig. 2A) and remains broadly distributed by 15-somites (Fig. 2B). At 24 hpf, *ndrg2* is strongly expressed in the brain, retina and spinal cord (Fig. 2C,D). At 48 hpf, its expression expands to the pectoral fin buds, myotome and the heart, with basal levels observed throughout the embryo (Fig. 2E,F). *ndrg3a* is broadly expressed at the shield stage (Fig. 3A) and is observed in the head region and pronephric ducts at 15-somites (Fig. 3B). At 24 hpf (Fig. 3C,D), *ndrg3a* is also seen in pharyngeal pouches, pectoral fin buds and myotome. By 48 hpf, *ndrg3a* signal is detected in the brain and the spinal cord in addition to the thymus, pronephric ducts and associated corpuscles of Stannius (Fig. 3E,F). *ndrg3b* signal is not detected between shield stage and 24 hpf (Fig. G,I,J); however, at 48 hpf it is observed in the brain and at lower levels in pectoral fin buds (Fig. 3K,L). At shield stage *ndrg4* is expressed ubiquitously (Fig. 2G) but becomes enriched in somites by the 15-somites stage (Fig. 2H). At 24 hpf and 48 hpf (Fig. 2I-L), *ndrg4* transcript is detected in the brain, the spinal cord, the pectoral fin buds, the intermediate cell mass of the mesoderm, and posterior-most pronephric duct.

**Figure 1.**
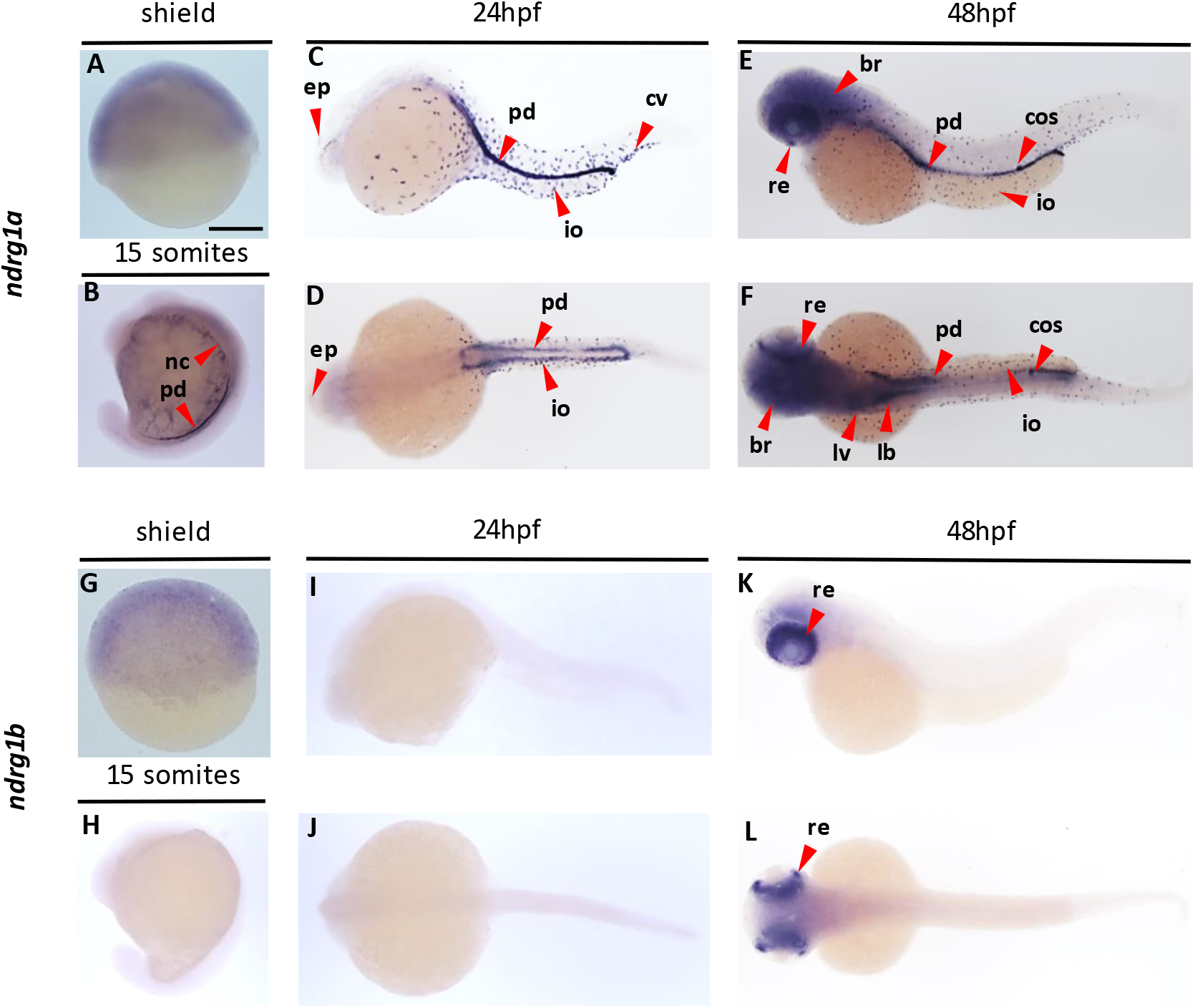
Gene expression analysis of *ndrg1*. WISH analysis revealing the distribution of *ndrg1a* (A-F) and *ndrg1b* (G-L) transcripts in zebrafish embryos at shield (A, G), 15 somites (B, H), 24 hpf (C, D, I, J) and 48 hpf (E, F, K, L) stages, imaged from lateral (A-C, E, G-I, K) and dorsal (D, F, J, L) views. Abbreviations: br (brain), cv (caudal vein), ep (epiphysis), cos (corpuscles of Stannius), ib (intestinal bulb), io (ionocyte), lv (liver), pd (pronephric ducts), nc (neural crest), and re (retina). Scale bar, 250 μm.

**Figure 2.**
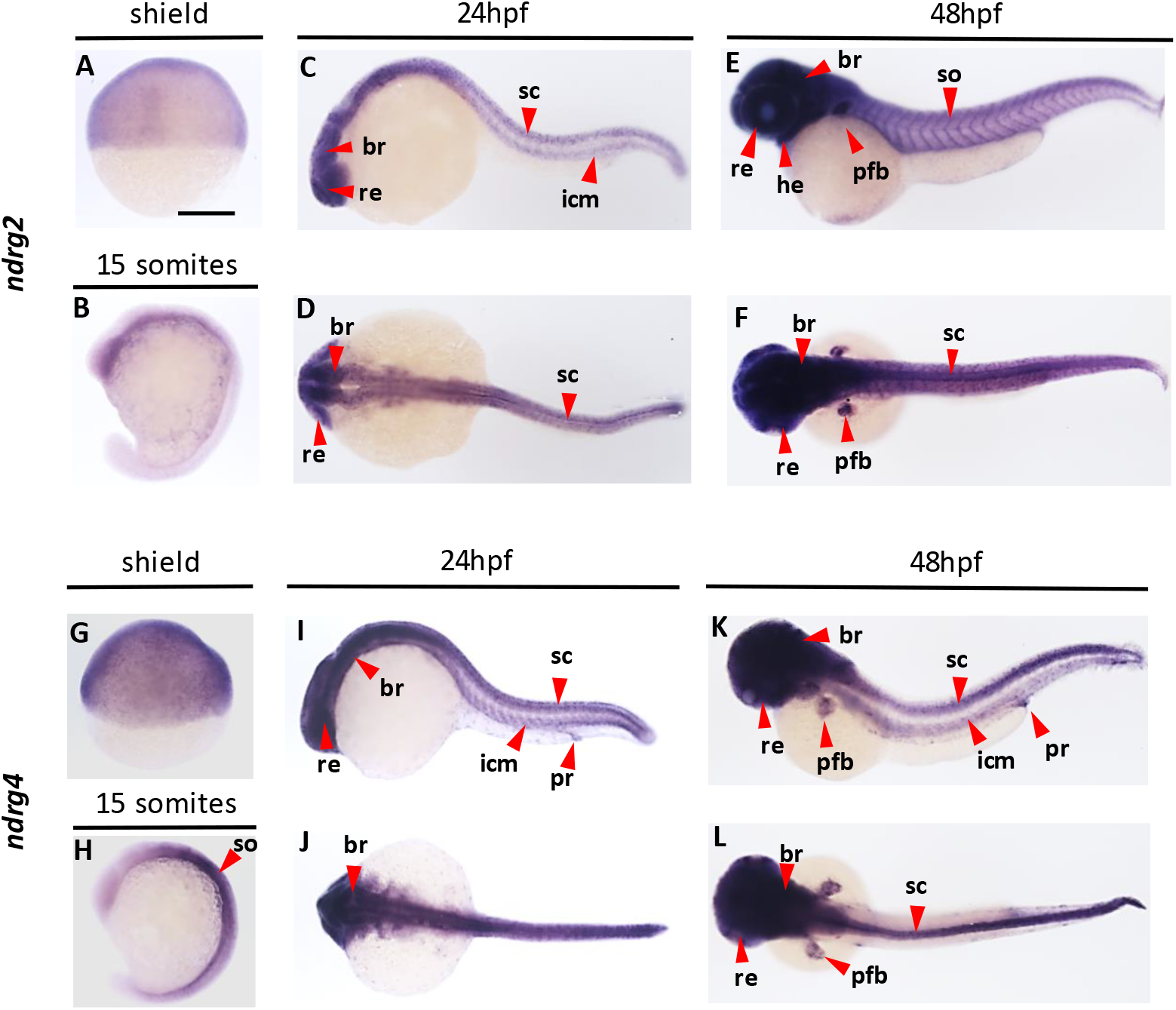
Gene expression analysis of *ndrg2* and *ndrg4.* WISH analysis revealing the distribution of *ndrg2* (A-F) and *ndrg4* (G-L) *transcripts* in zebrafish embryos at shield (A, G), 15 somites (B, H), 24 hpf (C, D, I, J) and 48 hpf (E, F, K, L) stages, imaged from lateral (A-C, E, G-I, K) and dorsal (D, F, J, L) views. Abbreviations: br (brain), he (heart), icm (intermediate cell mass of mesoderm), pfb (pectoral fin buds), pcv (posterior caudal vein), pr (proctodeum), re (retina), sc (spinal cord), and so (somites). Scale bar, 250 μm.

**Figure 3.**
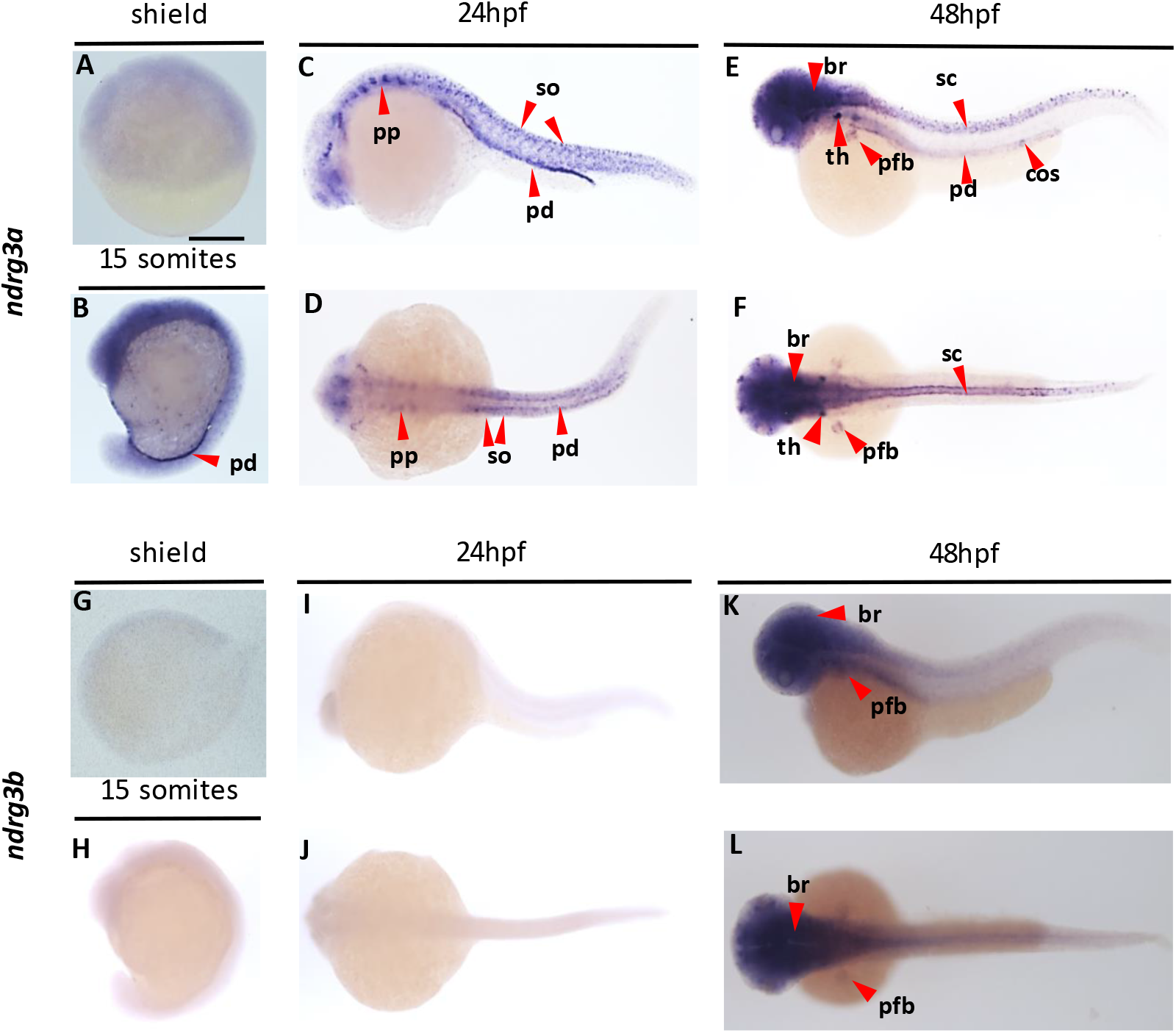
Gene expression analysis of *ndrg3.* WISH analysis revealing the distribution of *ndrg3a* (A-F) *and ndrg3b* (G-L) transcripts in zebrafish embryos at shield (A, G), 15 somites (B, H), 24 hpf (C, D, I, J) and 48 hpf (E, F, K, L) stages, imaged from lateral (A-C, E, G-I, K) and dorsal (D, F, J, L) views. Abbreviations: br (brain), cos (corpuscles of Stannius), pfb (pectoral fin buds), pp (pharyngeal pouches), pd (pronephric ducts), sc (spinal cord), so (somites), and th (thymus). Scale bar, 250 μm.

### Stage-matched embryos are appropriate controls for hypoxia-treated embryos

To gain an understanding of the transcriptional regulation of *ndrgs* in response to low O_2_, we exposed 24 hpf embryos to varying concentrations and durations of hypoxia and analyzed transcript levels using qPCR. Given that O2 deprivation delays or arrests zebrafish development, an important consideration for these experiments is the appropriate normoxic control group to use. We therefore began our analysis by considering control groups for embryos exposed to 4 or 8 hours of anoxia. Time zero normoxic controls are embryos that are the same age as the experimental group at the onset of treatment (24 hpf). Age-matched controls are the same age (hpf) as the experimental groups (28 hpf for embryos subjected to 4 hours of anoxia and 32 hpf for embryos exposed to 8 hours of anoxia). When using time zero controls, qPCR results revealed that transcript levels were significantly up-regulated for *ndrg1a* following 4 and 8 hours of anoxia (3 and 8-fold up-regulation, respectively), while other members of the *ndrg* family increased to a lesser extent (2-fold or less) (Fig. S1A). However, the use of age-matched controls resulted in a different outcome, with *ndrg1a* and *3a* being significantly up-regulated following 4 and 8 hours of anoxia while *ndrg2, 3b* and *4* were down-regulated and *ndrg1b* remained undetected (Fig. S1B). These differences in transcript levels using time zero and normoxic controls are most likely explained by dynamic gene expression during development, consistent with RNA Seq repository data (EMBL Zebrafish Expression Atlas) showing that *ndrg* expression levels change significantly between 24 and 48 hpf. These observations pointed to the importance of matching the developmental stage of the control and experimental groups, or, in other words, using stage-matched controls.

Several criteria were used to match the developmental stage of the experimental and control groups: the overall length of the embryo, the length of the yolk extension, head curvature, and level of pigmentation of the eye and the body (Fig. S2). Based on these criteria, the following normoxic control groups were selected for an expanded group of experimental conditions (hypoxia treatment of 24 hpf embryos/normoxic control group):4 h anoxia/26 hpf normoxic control (Fig. S2A and B), 8h of anoxia/27 hpf normoxic control (Fig. S2C and D), 4h 3% hypoxia/27 hpf normoxic control (Fig. S2E and F), 8h 3% hypoxia/30.5 hpf normoxic control (Fig. S2G and H).In addition to considerations about normoxic control groups, we sought for an appropriate reference gene to use for the qPCR analysis. A literature search identified translation elongation factor 1a *(ef1a)* as a strong candidate, whose levels are reported to remain constant under hypoxia [82, 83]. All experiments described below were performed using this reference gene and stage-matched controls.

### Differential regulation of members of the *ndrg* family in response to low oxygen

Cells adapt in distinct manners to varying levels of O_2_. Hypoxia (mild to severe) generally elicits metabolic reprogramming via HIF-1α-dependent transcriptional up-regulation of key genes that mediate the adaptive response [14, 15]. In contrast, anoxia-tolerance involves metabolic arrest, during which most ATP-demanding processes are suppressed, except for those that are essential for survival [16, 24]. To gain an understanding of the range of hypoxia conditions that elicit *ndrg* up-regulation and identify the members of this family that may promote hypoxia adaptation, we subjected embryos to hypoxia (3% O_2_) or anoxia (0% O_2_) for 4 or 8 hours.

In response to 4 hours of hypoxia, the transcript levels of none of the *ndrg* family members were significantly altered; *ndrg1b* expression was undetected (Fig. 4A). After 8 hours of hypoxia, *ndrg1a* was moderately up-regulated (1.9-fold). *ndrg1b* was also up-regulated (3-fold); however, this was not determined to be significant due to a large standard error. The expression of other members was not significantly altered (Fig. 4B).

**Figure 4.**
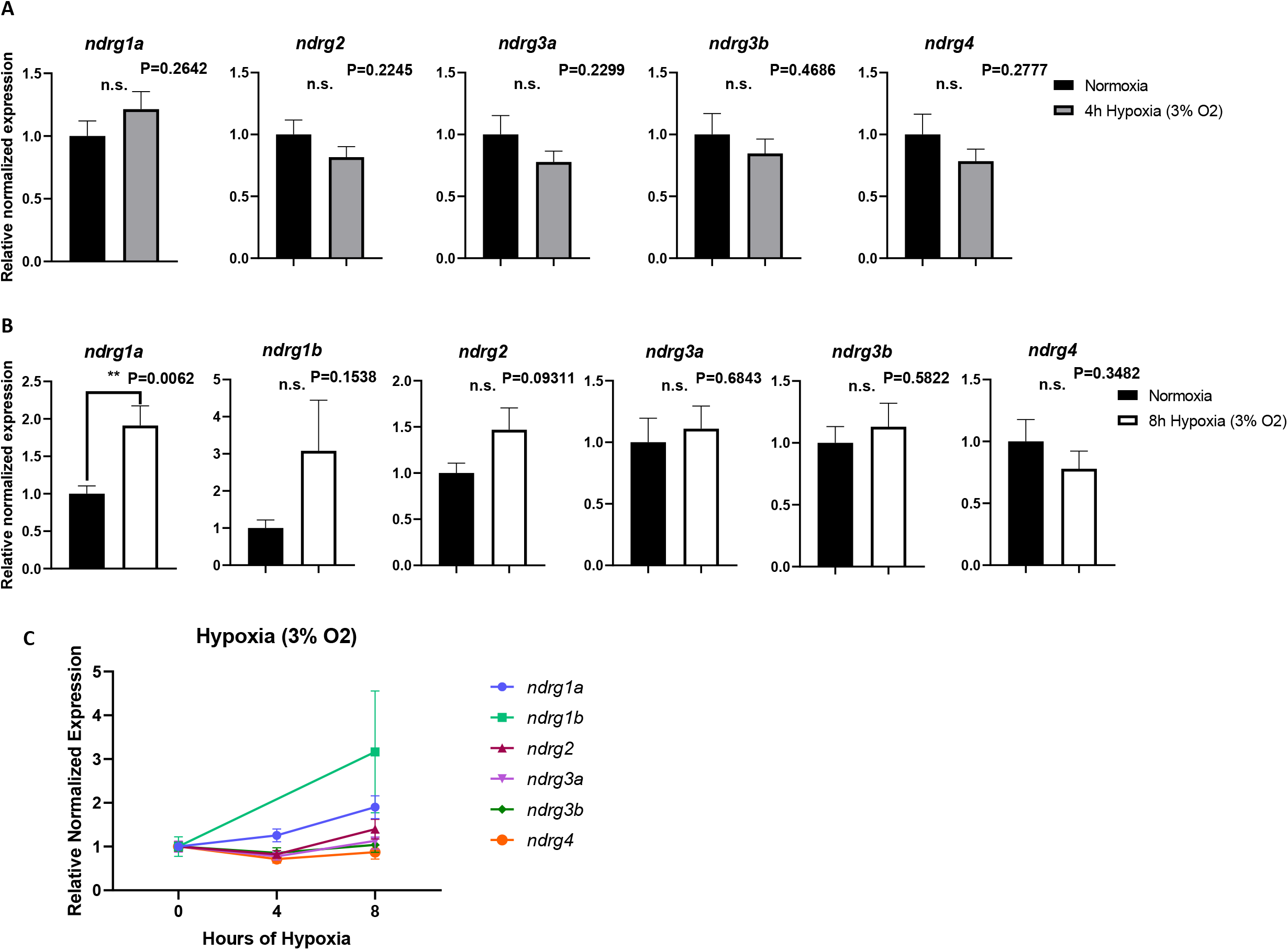
Changes in *ndrg* transcript levels in response to hypoxia (3% oxygen). (A, B) Real-time qPCR analysis of 24 hpf zebrafish embryos exposed to 4h (A, grey bars) or 8h (B, white bars) of hypoxia relative to normoxic (stage-matched) controls (A, B, black bars). (C) Linear summary of A and B. Expression levels were normalized to *ef1a.* The y-axis in the graphs represents the relative normalized expression of each gene. All fold changes were derived using the formula, 2 ^-(ΔΔ CT)^. Reactions were run in triplicate with 8 biological replicates (n=8). Bars represent standard error (SE). Unpaired, two-tailed t-test.

Following 4 hours of anoxia, *ndrg1a* was up-regulated ~ 2-fold. In contrast, *ndrg1b, ndrg3b,* and *ndrg4* were significantly down-regulated (~9-fold, ~2-fold, and ~2.9-fold, respectively), while *ndrg2* and *ndrg3a* expression was not significantly altered (Fig. 5A). By 8 hours of anoxia, *ndrg1a* was further up-regulated ~ 9-fold, and we also observed a slight up-regulation of *ndrg3a,* albeit less than 2-fold (1.9) (Fig. 5B). *ndrg2* (2-fold) and *ndrg4* (~1.67-fold) were down-regulated following 8 hours of anoxia and there was no significant change in the expression of *ndrg1b* (Fig. 5B).

**Figure 5.**
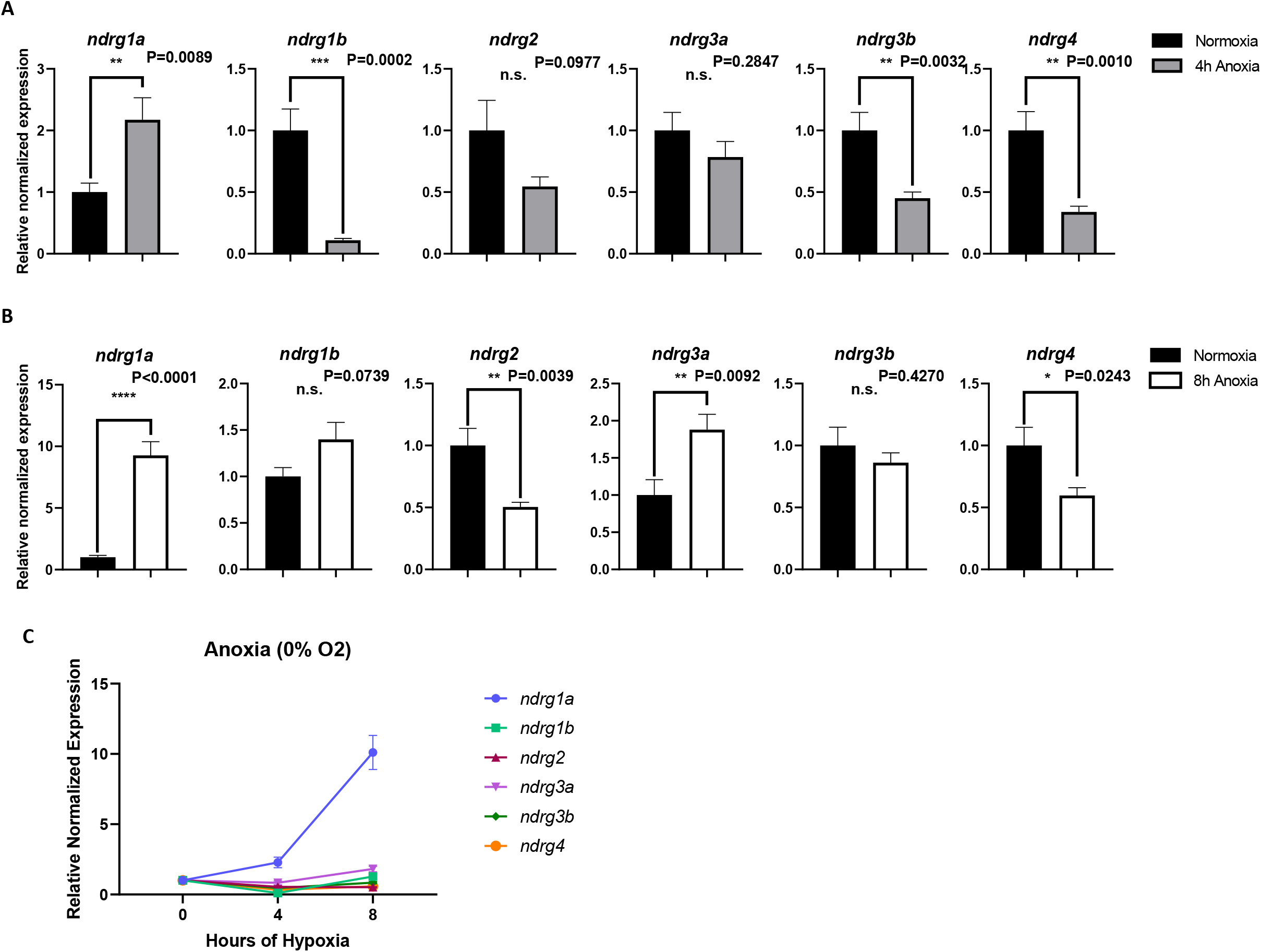
Changes in *ndrg* transcript levels in response to anoxia (0% oxygen). Real-time qPCR analysis of 24hpf zebrafish embryos exposed to 4h (A, grey bars) or 8h (B, white bars) of anoxia relative to normoxic (stage-matched) controls (A, B, black bars). (C) Linear summary of A and B. Expression levels were normalized to *ef1a.* The y-axis in the graphs represents the relative normalized expression of each gene. All fold changes were derived using the formula, 2 ^-(ΔΔ CT)^. Reactions were run in triplicate with 8 biological replicates (n=8). Bars represent standard error (SE). Unpaired, two-tailed t-test.

Overall, these data reveal that the *ndrg* family is differentially regulated in response to low O_2_. *ndrg1a* is the most hypoxia-responsive member of the family during early development and is transcriptionally up-regulated in response to severe and prolonged O_2_ deprivation. This finding is consistent with a previous study in cancer cells revealing that members of the NDRG family mediate long-term adaptation to hypoxia [67]. Despite the lack of reported HIF-1α binding sites in its promoter region, *ndrg3a* transcript levels also appear to be up-regulated under anoxia. Other members of the *ndrg* family may not be hypoxia-responsive at 24 hpf, at least not at the transcriptional level.

### Changes in the spatial distribution of *ndrg1a* in response to anoxia

To verify that *ndrg1a* is indeed hypoxia-responsive and determine if any changes in the spatial distribution of *ndrg1a* transcript occur following the most stringent hypoxia treatment, we performed WISH using 24 hpf embryos that were exposed to 8 hours of anoxia. Even though WISH is not a quantitative method to assess gene expression levels, we reasoned that the amount of transcript can at least be directly compared if control (stage-matched) and experimental (anoxia-treated) embryos were processed simultaneously during the color reaction step of WISH.

Following prolonged exposure to anoxia, *ndrg1a* transcript levels appear to be maintained at normoxic values in the pronephric duct and ionocytes but are noticeably elevated in the epiphysis (Fig. 6A,C). Interestingly, *ndrg1a* signal is also observed in tissues where this gene is not normally expressed (or expressed at levels that are below the detection limit) under normoxic conditions. Among these tissues, the head vasculature was prominently labeled in anoxia-treated embryos, namely: the primordial midbrain channel (pmbc), the primordial hindbrain channel (phbc), the mid-cerebral vein (mcb) and the aortic arch (aa) (Fig 6D, D’). *ndrg1a* transcript is also seen in the inner ear (otic vesicle, Fig. 6B, B’, D, D’) and in a segmentally-repeated pattern in the trunk (Fig. 6B, B’’, D, D’’) that apparently corresponds to somites. The mesoderm-expanded expression explains the thickened anterior-posteriorly oriented stripes of *ndrg1a* label observed from a dorsal view (compare Fig 6C and C’’ with D and D’’)

**Figure 6.**
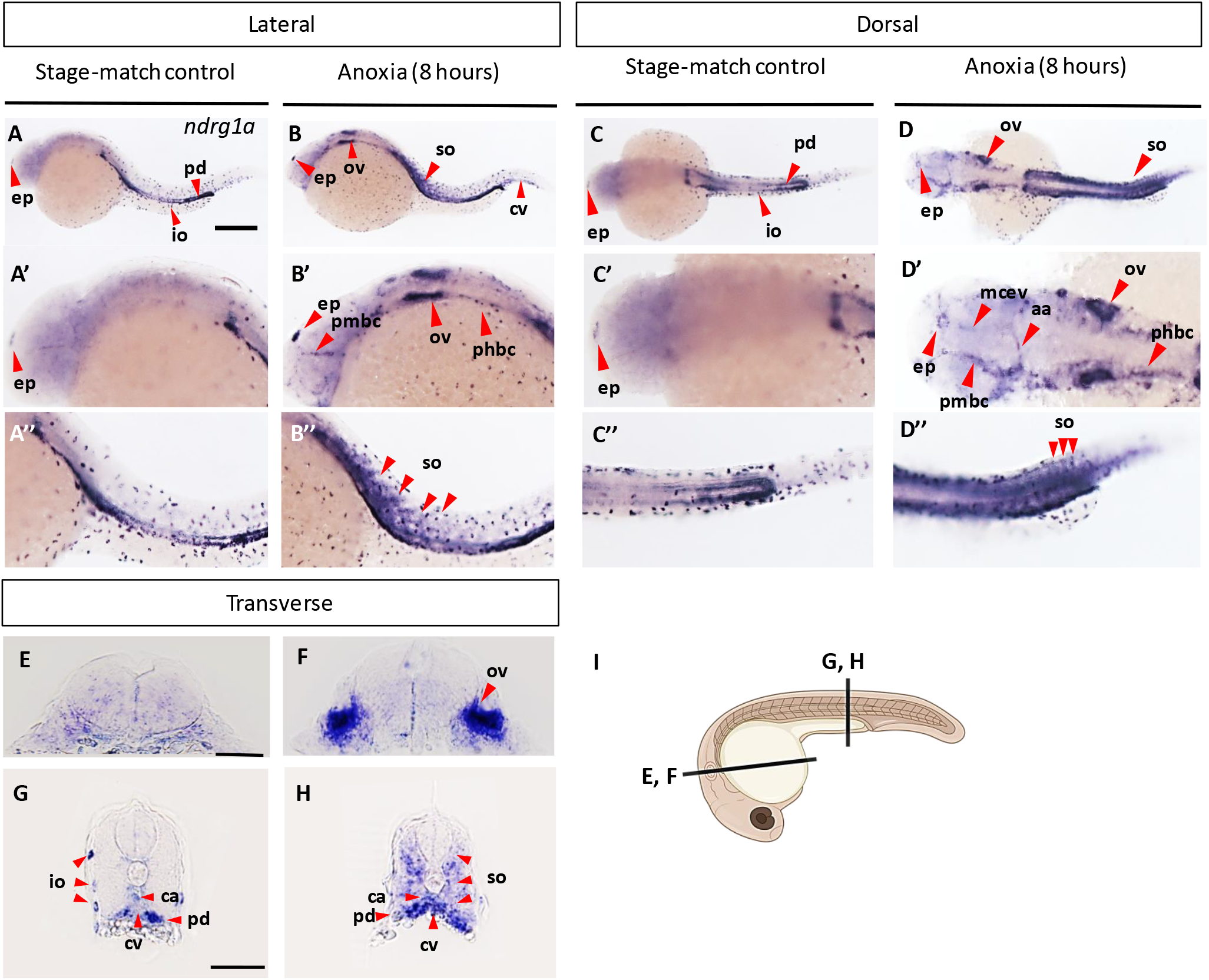
Analysis of *ndrg1a* expression in zebrafish embryos in response to 8h of anoxia. WISH analysis revealing the distribution of *ndrg1a* transcript in 24hpf zebrafish embryo exposed to 8h of anoxia (B-B’’, DD’’, F, H) relative to normoxic (stage-matched controls) (A-A’’, C-C’’, E, G) imaged from a lateral view (A-B’’), a dorsal view (C-D’’) and cross sectional (E-H) views through the otic vesicles (E,F) and the trunk (G,H), as indicated in panel I (illustration generated using biorender). (A’-D’’ are magnified views of (AD). Abbreviations: aa (aortic arch), cv (caudal vein), ep (epiphysis), io (ionocyte), mcev (mid-cerebral vein), ov (otic vesicle), pcv (posterior cardinal vein), phbc (primordial hindbrain channel), pmbc (primordial midbrain channel), and pd (pronephric ducts), and so (somites). Scale bar in A, 250 μm. Scale bar in E,G, 50 μm.

Cross sections of control and anoxia-treated embryos confirmed the anoxia-induced expression of *ndrg1a* in otic vesicles (Fig. 6F) and at basal levels throughout the somites, with some puncta of more intense label (Fig. 6G). In addition, the sections revealed elevated expression of *ndrg1a* in the caudal aorta and vein (Fig. 6G).

In summary, this analysis revealed that, following prolonged anoxia, *ndrg1a* transcript is maintained or expressed *de novo* in tissues in which it is present under normoxic conditions (epiphysis, pronephric duct and ionocytes) and expanded to additional tissues (vasculature, otic vesicles and somites). The overall increase in *ndrg1a* across multiple tissues accounts for the dramatic 9-fold up-regulation in transcript observed using q-PCR (Fig. 5B). These findings reveal hypoxia-dependent transcriptional regulation of *ndrg1a* in an intact, developing organism and identify tissues in which *ndrg1a* and other members of this family may play a protective role following severe and prolonged hypoxia.

## DISCUSSION

### Expression of *ndrg* family members during early development

WISH analysis of the *ndrg* family shows that during early development (shield), *ndrgs* are broadly expressed, with the exception of *ndrg3b* that is below detection levels (shield to 24 hpf). The overlapping expression of *ndrg1a, 1b, 2, 3a, and4* suggests that these genes may be functionally redundant. From mid-somitogenesis onward, *ndrgs* acquire more distinct spatial distribution patterns, consistent with previous studies revealing expression of members of this family in different cell types in the mouse brain [70] and organs/tissues of *Xenopus laevis* and *tropicalis* embryos [45, 68, 69].

Furthermore, the distribution of *ndrgs* is quite similar in fish and amphibian *(Xenopus tropicalis)* embryos. *ndrg1a* is observed in the eye, neural crest, pronephric duct, the intestinal bulb, and the liver of zebrafish and *Xenopus* embryos [45, 68, 69]. However, *ndrg1a* distribution in *Xenopus* appears broader than that of zebrafish, as it is also reported in the frog notochord, branchial arches, and pancreas. Similarly to zebrafish, the expression of *Xenopus ndrg2* is enriched in the nervous system; although there are also clear differences between these organisms since zebrafish *ndrg2* is prominent in somites while *Xenopus ndrg2* is found throughout the epidermis [45]. Zebrafish *ndrg3a* and *Xenopus ndrg3* are both present in the brain and spinal cord, but the former also localizes in the pharyngeal pouches, pronephric duct, thymus, and somites while the latter is enriched in the heart and otic vesicles [45]. *ndrg4* is strongly expressed throughout the nervous system in both zebrafish and *Xenopus,* but only observed in the zebrafish intermediate cell mass and the *Xenopus* pronephric duct [45]. Overall, these expression patterns suggest that the function of Ndrgs is at least partially conserved between fish and amphibians.

### *ndrgs* respond differentially to hypoxia

While transcription is an energy-demanding process that is suppressed under severe hypoxia [17], genes that mediate hypoxia adaptation are generally up-regulated [14, 15]. Our qPCR data reveal that among the *ndrg* family members, *ndrg1a* is the most hypoxia-responsive and that prolonged (8 hours) anoxia elicits the strongest increase in transcript levels. These findings corroborate with data from previous studies revealing that zebrafish and mammalian NDRG1 have HREs in their promoter region and are up-regulated in a HIF-1α-dependent manner in response to hypoxia [58, 59, 84]. *ndrg3a* is also up-regulated under anoxic conditions, albeit to a lesser extent than *ndrg1a* and does not have confirmed HREs, suggesting that other transcription factors contribute to its up-regulation.

Our qPCR data also revealed that *ndrg2, 3b,* and *4* are either unchanged or down-regulated, which can be explained in several ways. Unchanged values may reflect that the transcripts are stabilized, as previously reported for other genes [85], or that the rates of synthesis and degradation are equally matched. Downregulation could be due to mRNA decay exceeding the rate of synthesis or active repression of gene expression to conserve ATP [16, 17, 85, 86]. However, repression seems unlikely, as it is generally reserved for genes whose protein products are required for energetically-demanding processes (e.g. *elongation factor 5A* that mediates translation) [17]. Given that HREs have been reported in zebrafish *ndrg1a, 1b* and human *NDRG2* regulatory regions [58, 84], it is surprising that our qPCR analysis revealed that transcript levels of *ndrg1b* and *2* are either unchanged or decreased under low O_2_. With regards to *ndrg1b,* the spatially-restricted distribution of this transcript may account for the increase with low p-value we observed. It is also possible that a milder hypoxia treatment may be required to elicit up-regulation of *ndrg1b* and *ndrg2.* Indeed, a previous study revealed that hypoxia (5%) but not anoxia exposure of 24 hpf zebrafish embryos, caused the up-regulation of *igfbp-1, epo* and *vegf* [83]. Another explanation is that hypoxia-induced transcriptional regulation is dynamic, and additional developmental stages should have been tested to capture up-regulation of these genes, as was previously shown for *igfbp-1 and vegf* that are up-regulated in hypoxia-exposed 36 hpf, but not 24 hpf embryos [83].

These data reveal that members of the Ndrg family are differentially regulated in response to hypoxia and that *ndrg1a/ndrg3a* may be implicated in long-term adaptation to severe hypoxia.

### *ndrg1a* is up-regulated in metabolically-demanding tissues following prolonged anoxia

Previous studies using human cancer cells [58, 59] or homogenous cell lines (trophoblasts) [87] have revealed that *NDRG1* is up-regulated in response to hypoxia. However, little is known about how this response is orchestrated across multiple tissues of a whole organism. Using WISH, we investigated the distribution of *ndrg1a* in 24 hpf zebrafish embryos exposed to 8 hours of anoxia, a treatment that elicits the most robust increase in *ndrg1a* transcript. Given that these conditions are very stringent, we reasoned that any tissue/organ in which *ndrg1a* levels are significantly increased must require the activity of this protein to adapt to low O_2_.

Our study revealed that, following anoxia, *ndrg1a* is up-regulated in the epiphysis and possibly the pronephric duct, ionocytes, and endodermal organs (although the WISH procedure was not sensitive enough to detect an increase relative to the already high normoxic levels of *ndrg1a* in these cells). The epiphysis, also known as the pineal gland, receives information about the light-dark cycle from the environment and produces the hormone melatonin in response to this information. Melatonin has multiple cellular functions, including reducing oxidative stress [88], which is elevated under hypoxia and can cause cell death [89]. In addition to responding to light-dark stimuli, the pineal is also hypoxia-responsive, as stabilized Hif-1α modulates clock gene expression in zebrafish pineal cells [90, 91]. Given that *ndrg1a* is a Hif-1α target, it is possible that Ndrg1a is implicated in the regulation of clock genes and melatonin production under low O_2_ [92]. The liver is quite effective at taking up O_2_ and is normally well supplied by the bloodstream; nevertheless, it is susceptible to hypoxic injury and associated complications [93]. In contrast, the intestine normally experiences wide fluctuations in O_2_ throughout the day with some regions becoming hypoxic. Genes that aid in the maintenance of the hypoxic intestine are HIF-1α-regulated, providing a potential explanation for the expression of *ndrg1a* in this tissue [94, 95]. The function of *ndrg1a* in the pronephric duct and ionocytes is unclear, but these cells rely on the metabolically-demanding sodium-potassium ATPase pump to maintain ionic gradients and hence are likely to be sensitive to O_2_ depletion [96].

In addition to enhanced expression of *ndrg1a* in the epiphysis, we also observed expansion of *ndrg1a* distribution to tissues/organs where it is not present under normoxic conditions, namely the inner ear (otic vesicles), head vasculature, and somites. Previous studies have demonstrated that mutations in *NDRG1* are associated with Charcot-Marie-Tooth disease type 4D (CMT4D) [80], a demyelinating neuropathy that causes hearing loss in humans. Furthermore, hypoxia can cause hearing loss [97–100]; thus it is possible that Ndrg1a protects the inner ear or/and connected auditory nerve fibers from hypoxia-induced damage. Vascular sprouting is a well-documented hypoxic response to maximize O_2_ delivery [101]. NDRG1 was previously shown to mediate endothelial cell migration under intermittent hypoxia [102], raising the question of whether its upregulation under anoxia serves a similar purpose in head vasculature. Somites give rise to skeletal muscle cells, which experience cellular hypoxia and lactic acidosis during exercise that is further exacerbated by environmental hypoxia. It is possible that Ndrg1a protects muscle cells from acidosis or promotes hypometabolism in these cells.

Even though other members of the *ndrg* family are not transcriptionally up-regulated under anoxia (or at least not as significantly as *ndrg1a*), there is evidence that they can be post-translationally modified in response to hypoxia [67, 103–105]. In this regard, it is interesting that *ndrg2, 3a, 3b and 4* are expressed in the pectoral fin buds, which are known to play a respiratory role in fish [106, 107]. Furthermore, these genes are expressed in several metabolically-demanding tissues, including the brain, spinal cord, heart, and kidney.

In summary, we have shown that *ndrgs* are distributed across a range of hypoxia-sensitive/responsive tissues and that the levels of *ndrg1a* and *3a* are selectively increased following prolonged exposure to anoxia. Future studies will address whether members of this family promote hypoxia adaptation of the tissues and organs in which they are expressed.

## Supporting information

Hypoxia Regulation of ndrgs Supplemental Figures

## ACKNOWLEDGMENTS

We would like to acknowledge J. Park and B. Weinstein for their helpful comments on our manuscript. We would like to acknowledge T. deCarvalho and S. Larson of the Keith Porter Imaging Facility (KPIF, University of Maryland Baltimore Country) for obtaining images during times of COVID-19-related limited access to KPIF. This project was supported by funding from the Department of Defense (W81XWH-16-1-0466) and the National Institute of Health/NICHD (R21HD089476) to R. Brewster. N. Le was supported by a National Institute of Health /NIGMS MARC U*STAR (T34 HHS 00001) National Research Service Award to UMBC. T. Hufford was supported by CBI (NIGMS/NIH T32 GM066706) and IMSD (NIGMS/NIH T32 GM055036) training grants.

## AUTHOR CONTRIBUTIONS

N. Le executed the WISH and qPCR experiments and analyzed the data; T. Hufford performed qPCR experiments, plotted and analyzed the qPCR data and R. Brewster planned and oversaw the project and analyzed the data; N. Le wrote the first draft of the manuscript, T. Hufford and R. Brewster edited the manuscript.

## NONSTANDARD ABBREVIATIONS

NDRG: N-myc downstream regulated gene
HIF-1α: Hypoxia-inducible factor 1 alpha
HRE: Hypoxia response element
PHD2: Prolyl hydroxylase domain protein 2
VHL: von Hippel-Lindau protein
EPO: Erythropoietin
VEGF: Vascular endothelial growth factor
CREB: cAMP-response element binding protein
Myc: Myelocytomatosis proto-oncogene, basic helix-loop-helix transcription factor
NF-kB: Nuclear factor kappa-light-chain-enhancer of activated B cells
STAT: Signal transducer and activator of transcription
Drg1: Downregulated gene 1
Cap43: Calcium associated protein 43 kDa
Rit42: Reduced in tumor, 42 kDa
RTP: Reducing agents and tunicamycin-responsive protein
PROXY-1: Protein regulated by oxygen 1
AP-1/2: Activator proteins 1 & 2
LAMP1: Lysosomal-associated membrane protein 1
Rab4: Ras-related protein Rab-4A
TNF-α: Tumor necrosis factor alpha
WISH: Whole-mount *in situ* hybridization
Igfbp-1: insulin-like growth factor-binding protein 1

