## Supplementary figures and images for "Hypoxia Regulation of *ndrgs*"

### Hypoxia Regulation of ndrgs Supplemental Figures

#### Slide 1
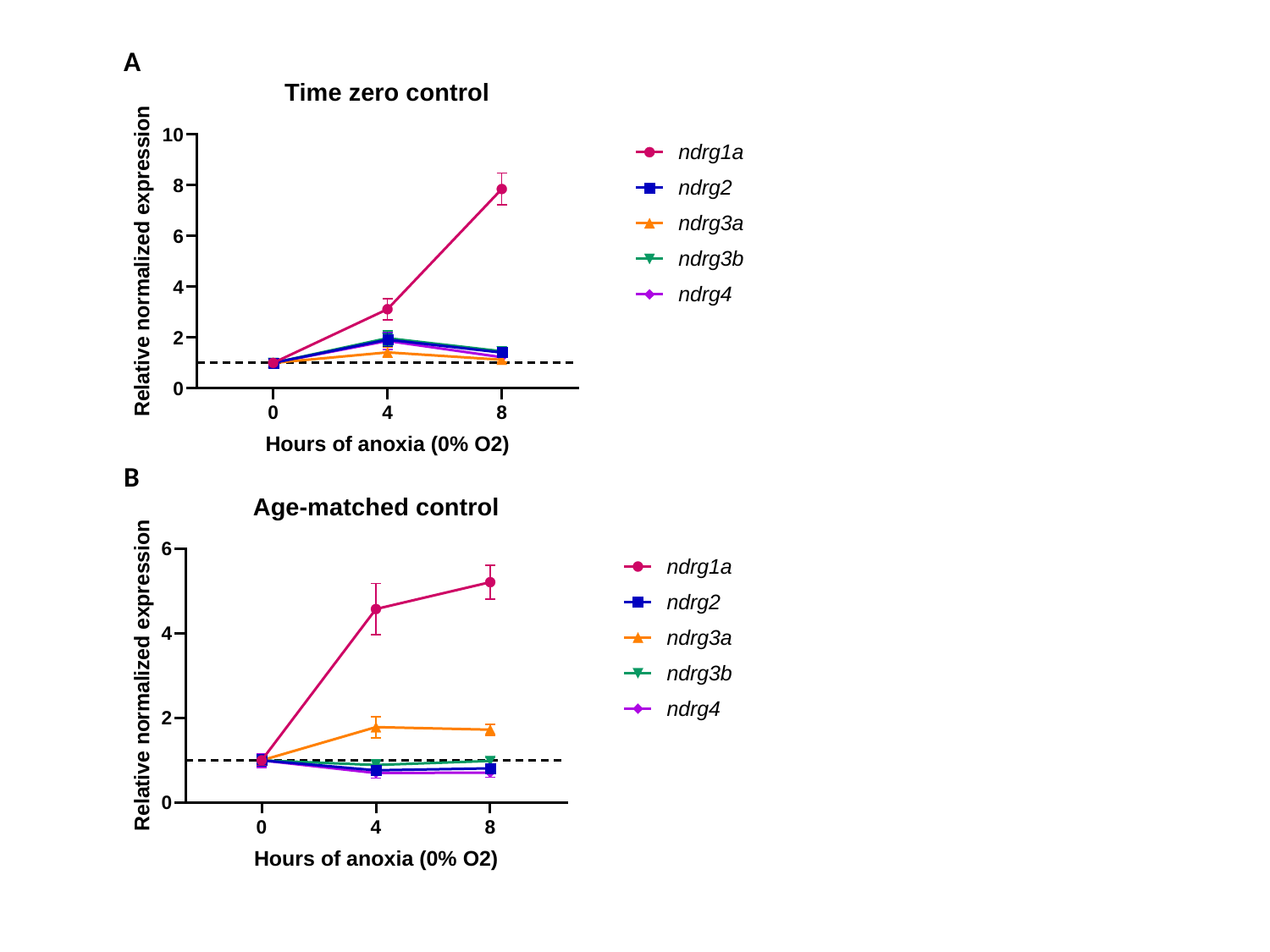

A
B

#### Slide 2
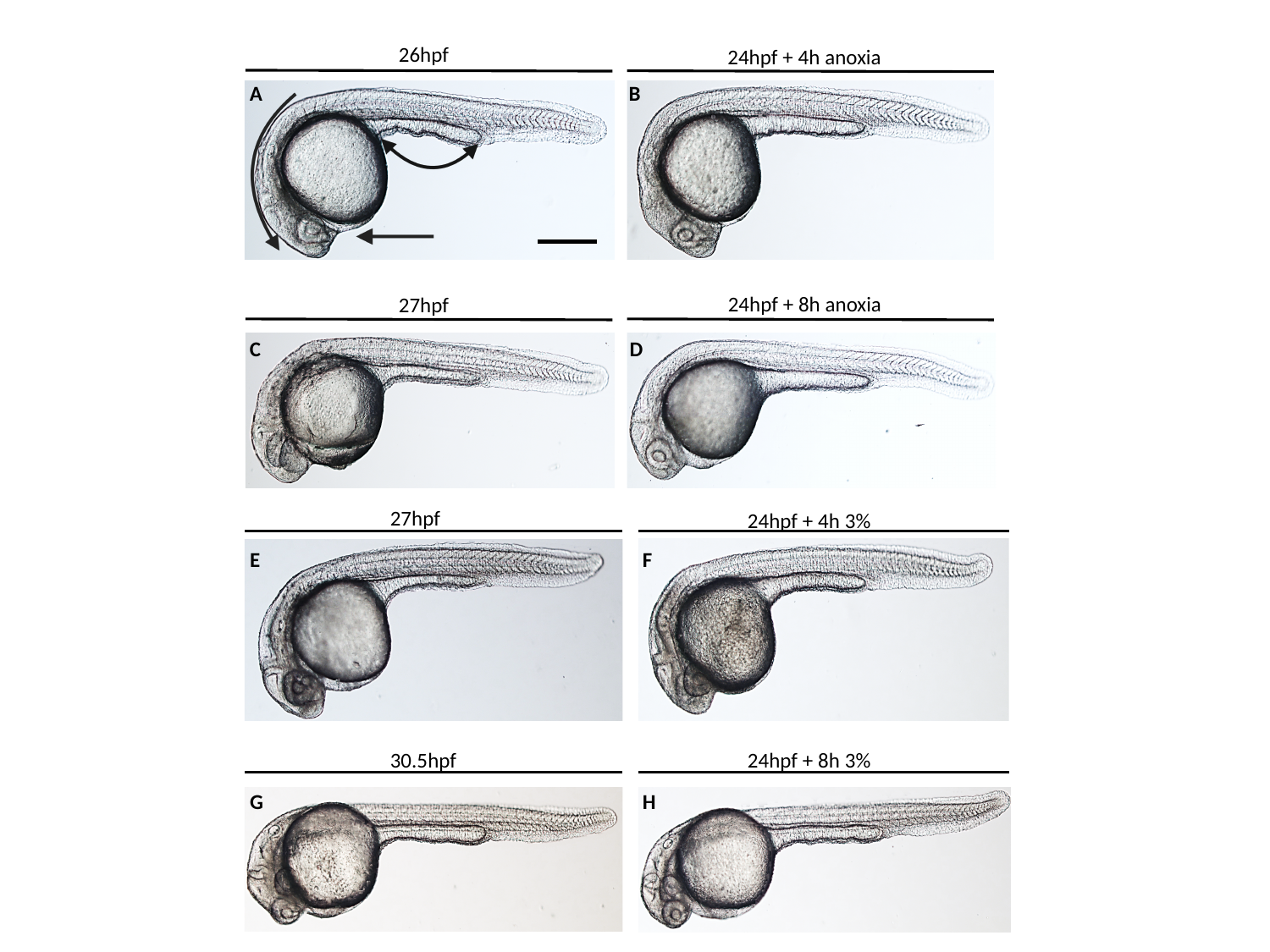

26hpf
24hpf + 4h anoxia
A
B
24hpf + 8h anoxia
27hpf
C
D
27hpf
24hpf + 4h 3%
E
F
24hpf + 8h 3%
30.5hpf
G
H
